# Obesity promotes conserved inflammatory and metabolic transcriptional programs in mouse and human colon tumors

**DOI:** 10.1101/2025.06.19.660576

**Authors:** Elaine M. Glenny, Tengda Lin, Victoria M. Bandera, Babak Mirminachi, Saratchandra S. Khumukcham, Biljana Gigic, Christy A. Warby, Olena Aksonova, Michael F. Coleman, Alessandro Carpanese, Calista Busch, Caroline Himbert, Jennifer Ose, David A. Nix, Kenneth Boucher, Peter Schirmacher, Ildiko Strehli, Sheetal Hardikar, Jessica N. Cohan, Jolanta Jedrzkiewicz, Alexander Brobeil, Martin A. Schneider, Christoph Kahlert, Erin M. Siegel, Doratha A. Byrd, Adetunji T. Toriola, David Shibata, Christopher I. Li, Jane C. Figueiredo, Aik Choon Tan, Jatin Roper, Cornelia M. Ulrich, Stephen D. Hursting

## Abstract

**Background:** The global prevalence of obesity, an established risk and progression factor for colon cancer, is high and rising. Unfortunately, the mechanisms underlying the obesity-colon cancer association are incompletely understood, and new molecular targets enabling more effective intervention strategies to break the obesity-colon cancer link are urgently needed.

**Objective:** This study integrated RNA sequencing data from mouse and human colon tumor samples, as well as human adipose samples, to rigorously establish obesity-associated transcriptomic signatures conserved between the two species.

**Methods:** We employed a mouse colon cancer model with colonoscopy-guided orthotopic transplantation of syngeneic *Apc-null;KrasG12D/+;Trp53-null;Smad4-null;tdTomato* colon tumor organoids. Epithelial cell adhesion molecule (EpCAM)-positive cells from murine tumors, and 193 human colon tumors and 188 human mesenteric adipose tissue samples from the ColoCare cohort underwent transcriptomic analyses.

**Results:** Diet-induced obesity reduced survival in the mouse model of colon cancer. Integrated transcriptomic analyses of EpCAM-positive murine tumor cells and bulk human tumors revealed obesity-driven enrichment of inflammation and metabolic pathways, including upregulation of genes involved in innate immune sensing (*TLR2*, *MYD88*, *IRF4*) and tumor microenvironment remodeling (*MMP9*, *TGFB1*, *SERPINE1*). Analysis of paired mesenteric visceral adipose tissue and tumor samples from the ColoCare cohort indicated that obesity amplifies inflammatory signaling pathways through unique adipose ligand-tumor receptor interactions.

**Conclusions:** These results establish obesity-associated adipose tissue dysregulation as a key inter-tissue modulator of biology, with concordant cross-species effects on tumor cell-intrinsic inflammatory and metabolic programs.

## INTRODUCTION

Colorectal cancer is the third most common cancer and the second leading cause of cancer deaths worldwide with approximately 1.9 million new cases and 900,000 deaths in 2022 and rising rates of early-onset colorectal cancer (1–3). Obesity, a metabolic disease defined as body mass index (BMI) ≥30 kg/m^2^, is a major risk and prognostic factor for colon cancer and at least 12 other cancer types (4). Patients with colon cancer and obesity have reduced overall survival and disease-free survival risk relative to patients with a normal BMI (<25 kg/m^2^) (5).

Obesity rates have tripled worldwide since 1975, and >70% of adults in the US are either overweight or obese (6, 7). However, critical biological pathways underpinning the obesity-colon cancer relationship remain incompletely understood, hampering efforts to develop effective mechanism-based nutritional or pharmacologic interventions to reduce the burden of obesity-driven colon cancer (8).

Expanded and dysregulated visceral adipose tissue (VAT) is a hallmark of obesity (8–11). In colon cancer, mesenteric VAT envelopes the colonic serosa and interfaces with the tumor vasculature, facilitating exchange of adipose-derived factors. Beyond its role as an energy storage site, VAT serves as an active endocrine tissue comprised of many cell types including adipocytes, immune cells, and fibroblasts that collectively secrete proteins and metabolites involved in the regulation of energy homeostasis and metabolic and inflammatory pathways (8, 11). Altered morphological and functional changes in VAT, as well as levels of several VAT-derived factors (particularly adipokines, cytokines, growth factors, and lipid mediators) may contribute to the enhancing effects of obesity on colon cancer risk and progression (8, 9, 11). However, actionable targets to disrupt the crosstalk between obesity-associated dysregulated VAT and the colon tumor have yet to be systematically identified and characterized.

Murine models are instrumental for deeply interrogating the mechanistic underpinnings of cancer, but they do not always recapitulate the biological complexities of human disease (12–15). Conversely, observational studies using human samples are invaluable for identifying candidate genes and pathways highly relevant to cancer development and progression but are typically limited to identifying associations rather than establishing causal relationships (16).

When evaluated in parallel, however, preclinical and human data have synergistically informed each other to accelerate the pace of translational research on obesity and cancer (17–20).

Herein we leveraged transcriptomic data from both preclinical and human studies to identify putative molecular targets for disrupting the obesity-colon cancer link. Specifically, through an integrated genomics analysis using a state-of-the-art obesity-driven orthotopic colon cancer mouse model, and paired tumor and mesenteric VAT samples from an international cohort of patients with newly diagnosed colon cancer (21), we identify obesity-responsive pathways in tumors, interactions between dysregulated VAT and the colon tumor, and procancer gene candidates conserved between species. These analyses disclose new molecular targets for disrupting obesity-driven colon cancer that are conserved across two species and reveal VAT-to-tumor paracrine signaling as a putative driver of inflammatory processes in the tumor.

## METHODS

### Study approval

All human study participants provided written informed consent and the research was approved by the Institutional Review Boards of all participating institutions. All mouse studies were approved by the Institutional Animal Care and Use Committee (IACUC) at Duke University.

### Study population

The study population was drawn from the international prospective ColoCare Study cohort (U01CA206110; Clinicaltrials.gov Identifier: NCT02328677) (21). This multicenter cohort study recruits male and female adult patients (18-89 years of age) with primary invasive colorectal cancer who were scheduled to undergo surgical intervention at participating research centers. Patients with stage I-III colon cancer with available colon tumor tissue were eligible for the current study. 193 male and female patients, of which 188 patients also had available fresh-frozen tumor-adjacent VAT, from the Huntsman Cancer Institute, Moffitt Cancer Center, University of Tennessee Health Science Center, and the Heidelberg University Hospital, Germany, were included in these analyses (**Table 1**). None of the patients had undergone neoadjuvant chemotherapy and only one patient had a BMI <18.5 kg/m^2^.

**Table 1.** Baseline clinicodemographic characteristics of patients with colon cance

|  | **Normal**<br>**(N=45)** | **Overweight**<br>**(N=82)** | **Obese**<br>**(N=66)** | **Overall**<br>**(N=193)** |
| --- | --- | --- | --- | --- |
| **Age at diagnosis** |  |  |  |  |
| Mean ± SD | 64.7 ± 13.7 | 63.2 ± 13.4 | 62.0 ± 12.1 | 63.1 ± 13.0 |
| 95% CI | (60.6, 68.8) | (60.3, 66.2) | (59.0, 64.9) | (61.3, 65.0) |
| Median (Min, Max) | 68.0 (19.0, 86.0) | 64.5 (18.0, 86.0) | 62.5 (27.0, 84.0) | 64.0 (18.0, 86.0) |
| **Sex** |  |  |  |  |
| Female | 32 (71.1%) | 27 (32.9%) | 34 (51.5%) | 93 (48.2%) |
| Male | 13 (28.9%) | 55 (67.1%) | 32 (48.5%) | 100 (51.8%) |
| **Race** |  |  |  |  |
| White | 43 (95.6%) | 74 (90.2%) | 58 (87.9%) | 175 (90.7%) |
| African American | 0 (0%) | 6 (7.3%) | 5 (7.6%) | 11 (5.7%) |
| American Indian or Alaska Native/Asian/Pacific Islander | 0 (0%) | 2 (2.4%) | 3 (4.5%) | 5 (2.6%) |
| Other | 2 (4.4%) | 0 (0%) | 0 (0%) | 2 (1.0%) |
| **Pre-surgery BMI<sup>B</sup>** |  |  |  |  |
| Mean ± SD | 22.4 ± 1.6 | 27.4 ± 1.3 | 35.1 ± 5.5 | 28.9 ± 6.0 |
| 95% CI | (22.0, 22.9) | (27.1, 27.7) | (33.7, 36.4) | (28.0, 29.7) |
| Median (Min, Max) | 22.4 (18.0, 24.9) | 27.4 (25.1, 29.8) | 33.4 (30.0, 59.1) | 28.0 (18.0, 59.1) |
| **Tumor stage at diagnosis** |  |  |  |  |
| I | 9 (20.0%) | 13 (15.9%) | 4 (6.1%) | 26 (13.5%) |
| II | 15 (33.3%) | 35 (42.7%) | 29 (43.9%) | 79 (40.9%) |
| III | 21 (46.7%) | 34 (41.5%) | 33 (50.0%) | 88 (45.6%) |
| **Microsatellite Instability Status (MSI) testing** |  |  |  |  |
| Microsatellite Stable | 26 (66.7%) | 50 (72.5%) | 45 (78.9%) | 121 (73.3%) |
| Microsatellite Unstable | 13 (33.3%) | 19 (27.5%) | 12 (21.1%) | 44 (26.7%) |
| Missing | 6 | 13 | 9 | 28 |

<sup>A</sup>Based on 193 patients with colon cancer with tumor transcriptomes available; 188 of these patients also provided visceral-adipose-tissue (VAT). n=89 from Huntsman Cancer Institute; n=65 from Heidelberg University Hospital; n=20 from University of Tennessee Health Science Center; n=19 from Moffitt Cancer Center.
<sup>B</sup>Body mass index (BMI) at cancer diagnosis or at enrollment.

### Animals and diets

Female C57BL/6Hsd mice were purchased from Inotiv (#044; West Lafayette, Indiana) and housed in a temperature- and humidity-controlled specific-pathogen-free facility with a 12- hour light cycle. Cages contained ALPHA-dri^TM^ bedding (Shephard Specialty Papers, Watertown, TN) and mice were randomized to be provided with ad libitum access to either a 10 kcal% fat purified diet (control diet; D12450J; Research Diets, Inc, New Brunswick, NJ) or a 60 kcal% fat purified diet (DIO diet; D12492; Research Diets, Inc) for duration of studies. At study termination, mice were anesthetized using 500 mg/kg 2,2-tribromoethanol and euthanized via thoracotomy when unresponsive via toe pinch.

### Organoid orthotopic transplantation

Dissociated *Apc-null;KrasG12D/+;Trp53-null;Smad4-null;tdTomato* (i.e., AKPST) organoids (a kind gift from Ömer H. Yilmaz, Massachusetts Institute of Technology) were mixed with Matrigel (#356231; Corning, NY) at a 1:3 ratio. 7-8 20 µl domes were plated in a 6-well plate and incubated at 37°C for 10-15 min. Following Matrigel polymerization, organoid media (Advanced DMEM-F12 media [Gibco, Carlsbad, CA] supplemented with 5% fetal bovine serum [Cytiva, Marlborough, MA], 2% B27 [Gibco], 1% L-glutamine [Gibco], and 1% antibiotic-antimycotic [#Gibco]) was added and cultures were maintained at 37 °C with 5% CO_2_.

Organoids were passaged every 2-3 days using cell recovery solution (Corning) to dissolve the Matrigel and TrypLE express enzyme (Gibco) to create a single-cell suspension before re- seeding.

AKPST organoids were orthotopically transplanted as previously described for other colon cancer organoid lines (22, 23). Briefly, Matrigel was chemically dissociated using 500 µl cell recovery solution for 30 min on ice. Organoids were pelleted, washed with cold PBS, and resuspended in 10% Matrigel with organoid media containing no FBS at a concentration of 10 organoids/µl. Mice were anesthetized with 1- 2% isoflurane, placed in the supine position, and kept warm on a heating pad for the procedure (SomnoSuite, Kent Scientific). Colons were flushed with water using a gavage needle to expunge fecal matter. Mice were injected with a single 70 µl bolus of AKPST organoids into the colonic submucosa via colonoscopy-guided injection (Coloview, Karl Storz, Tuttlingen, Germany) using a 16 in, 45-degree bevel, 33-gauge needle (Hamilton, Reno, NV).

### Survival analysis

Eight-week-old Control and DIO female C57BL/6Hsd mice were maintained on either the 10 kcal% fat control diet or 60 kcal% fat DIO diet, respectively, for 35 weeks to promote excessive weight gain and VAT accumulation in DIO mice. All mice were weighed immediately prior to orthotopic transplantation of 700 AKPST organoids in the colonic submucosa. Mice were weighed in a random order and survival was defined as the number of days post orthotopic transplant that mice were alive until >25% unintentional weight loss occurred.

### EpCAM+ enrichment from orthotopic colon tumors

Tumors were collected four weeks after organoid transplant, cut into 2-4 mm pieces, and stored on ice in Leibovitz’s L-15 media (Corning). Tumors were dissociated into single-cell suspensions using a mouse tumor dissociation kit (Miltenyi Biotec, Bergisch Gladbach, Germany) and a gentleMACS dissociator (Miltenyi Biotec). Resulting suspensions were sequentially filtered using 70 µm and 30 µm filters, centrifuged, and resuspended in RPMI 1640 media (Gibco). CD45+ cells were removed using CD45 MicroBeads (Miltenyi Biotec) and EpCAM+ cells were subsequently enriched from the CD45- fraction using CD326 (EpCAM) MicroBeads (Miltenyi Biotec) per manufacturer’s instructions. Cell fractions were centrifuged and pelleted cells were lysed with lysis buffer (Qiagen, Hilden, Germany) and stored at -80°C.

### RNAseq analysis for murine tumors

RNA was isolated from EpCAM+-enriched cell populations using the RNeasy micro kit (Qiagen). RNA was quantified on a Qubit 2.0 fluorometer (Life Technologies, Carlsbad, CA) and integrity was assessed on a TapeStation 4200 (Agilent Technologies, Santa Clara, CA). ERCC RNA spike-in mix (Thermo Fisher Scientific) was added to normalize total RNA. RNA sequencing libraries were prepared using a NEBNext Ultra II RNA library prep kit (New England Biolabs, Ipswitch, MA). mRNAs were enriched, fragmented, and used for cDNA synthesis. Prior to sequencing, cDNA fragments were end-repaired, adenylated, and ligated with universal adapters, and libraries were quantified by a Qubit 2.0 fluorometer and validated on a TapeStation 4200. Sequencing was performed on Illumina HiSeq instrument using a 2 x 150 bp paired-end configuration at a depth of >25 million reads per sample.

Raw data were converted to fastq files and de-multiplexed using Illumina’s bcl2fastq 2.17 software, allowing one mismatch for index sequence identification. Adapters were trimmed and low-quality reads were filtered using Trimmomatic (24). Spliced Transcripts Alignment to a Reference (STAR) aligned reads to the reference GRCm38 mouse genome and produced a counts matrix (25). Low abundance genes (<10 read counts in ≥7 samples) were removed from all downstream analyses and a normal shrinkage estimator was applied to log2 fold change (log2FC) estimates to enhance gene stability, particularly for genes with low counts and/or high variability (26). Principal coordinate analysis (PCA) plots were generated using the top 500 most differentially expressed genes based on FDR q-values based on DESeq2 analysis using the FactoMineR package (v2.8) (26, 27).

### RNAseq analysis for human VAT and tumors

Human colon tumor tissue and proximal VAT samples (within 1-3 cm from tumor) were collected and stored at -80 °C. All shipped samples were initially stored at -120 °C and shipped on dry ice to the Huntsman Cancer Institute (HCI). RNA from ∼25 mg of tumor tissue was extracted by the Biorepository and Molecular Pathology Shared Resource at the HCI using the Qiagen RNeasy Plus Mini Kit (Qiagen, Hilden Germany) per manufacturer’s protocol. RNA from ∼65 mg of VAT was extracted by bead beating in 1 ml TRIzol (Life Technologies, Carlsbad CA) and using the PureLink RNA Mini Kit (Invitrogen, Carlsbad CA) per manufacturer’s protocol with two modifications. All centrifugation steps were performed at 4 °C and an additional centrifugation step was added prior to the aqueous phase separation. RNA sequencing libraries were made using the NEBNext Ultra II Directional RNA Library Prep with rRNA Depletion Kit (New England Biolabs). Sequencing was performed at the High-Throughput Genomics at the HCI on either the Illumina NovaSeq 6000 or NovaSeq X instruments using a 2 x 150 bp paired-end configuration at a depth of ∼25 million reads per sample.

Raw sequencing data were converted to fastq files and analysis was performed using a customized in-house developed containerized RnaAlignQC workflow. This workflow removes adapter sequences from the paired fastq reads using CutAdapt, aligns the reads to GRCh38 using the transcript aware STAR aligner, estimates gene and isoform abundance with RNA-Seq by Expectation Maximization (RSEM), assigns uniquely mapping reads with featureCounts, and calculates a variety of QC metrics using FastQC, FastqScreen, Picard’s CollectRnaSeqMetrics, and MultiQC. Low abundance genes (<10 read counts in all samples) were removed.

Differential gene expression analysis was conducted separately for tumor (N=193) and VAT (N=198) samples using DESeq2 (v1.42.1)(26) in R(v4.3.3). We examined gene counts adjusted for relevant covariates [age (tertiles), sex (female/male), tumor stage (I/II/III), and study center (HCI/Moffitt Cancer Center/University of Tennessee Health Science Center/Heidelberg University Hospital)] using multiple contrasts of interest based on BMI (obese vs. non-obese, obese vs. normal, overweight vs. normal, and obese vs. normal). A normal shrinkage estimator was applied to log2 fold change estimates (26).

### Gene set enrichment analysis (GSEA)

GSEA (v4.3.3) was performed on DESeq2 normalized data using the MSigDB Hallmark gene sets (v2024.1) (28). Ranked gene lists were created using log2FC values generated by DESeq2 with the normal shrinkage estimator applied, and mouse genes were collapsed and remapped to human gene symbols. Default settings were used to generate a normalized enrichment statistic for all gene sets tested. Genes contributing to the leading edge in a subset of gene sets were visualized as chord diagrams using the circlize package in R (v0.4.16) (29). Custom gene sets were created by including all genes contributing to the leading edges of significantly enriched gene sets related to either metabolic or inflammatory processes for the identified BMI pairwise comparisons.

### VAT ligand-tumor receptor analysis

Ligand-receptor analysis was conducted on paired VAT and tumor samples from 188 ColoCare patients to investigate VAT-tumor signaling. Analysis was based on normalized counts, expressed as counts per million, to assess the correlation of gene expression for each ligand-receptor pair across VAT and tumor tissues (i.e., the correlation between a ligand in VAT and its cognate receptor in tumor). The analysis was conducted separately for samples from patients across the three BMI categories: normal, overweight, and obese. Ligand-receptor pairs were identified using three databases: FANTOM 5 (30), CellPhoneDB (31), and CellChat (32). A Pearson’s correlation threshold of >0.3 was set to determine significant ligand-receptor pairs, along with a requirement that at least one-third of samples from each BMI category were complete pairs (nonzero counts for both ligand and cognate receptor).

### Over-representation analysis

Over-representation analysis was performed using EnrichR (33) using 123 unique genes identified from VAT-ligand tumor-receptor pairs concordantly enriched (Pearson’s correlation coefficient >0.3) in samples from patients with colon cancer and obesity.

### Statistics

Statistical analyses were performed using GraphPad Prism (v10.2.2; GraphPad Software Inc., La Jolla, CA). Unpaired Student’s two-tailed t-tests were used to compare two groups. Survival analysis was conducted using log-rank (Mantel-Cox) test. p<0.05 was considered statistically significant. No samples were excluded from analysis.

For RNA-sequencing data, differential gene expression was determined using DESeq2(26) at a false discovery rate (FDR) q<0.05. Pairwise adonis test assessed for differences in gene expression patterns between groups where p<0.05 was considered significant. Analyses of MSigDB Hallmarks using GSEA or over-representation analysis were considered significant at FDR q<0.05 or adjusted p<0.05, respectively.

## RESULTS

### Diet-induced obesity reduces survival in an orthotopic murine organoid model of colon cancer

We determined the effect of diet-induced obesity (DIO) on survival in a colonoscopy-guided, organoid-based orthotopic transplantation model of murine colon cancer (22). In this model, the engraftment rate of *Apc-null;KrasG12D/+;Trp53-null;Smad4-null;tdTomato* (AKPST) organoids is >95% where successful transplantation is marked by tumor formation in the submucosal layer of the colonic wall (**Figure 1A-B**). Review by a veterinary pathologist demonstrated that tumors are high-grade, malignant adenocarcinomas with histological features, such as the presence of glands and the recruitment of multiple cell types including fibroblasts and diverse immune populations, that recapitulate human colon cancer (**Figure 1C-D**) (22). To generate Control (non-obese) and DIO groups, mice consumed either a low-fat control diet (i.e., 10% calories from fat) or high-fat DIO diet (i.e., 60% calories from fat), respectively, ad libitum for 36-38 weeks (**Figure 1E**), at which time DIO mice were >2-fold heavier than Controls (**Figure 1F**). To assess whether DIO accelerates obesity-associated colon cancer progression, we conducted a survival study in Control and DIO mice that were orthotopically transplanted with AKPST organoids (**Figure 1G**). The DIO group, relative to Controls, had reduced median survival (36.5 days vs. 56.5 days; hazard ratio 3.1; p=0.0002; **Figure 1H**). As expected, tumors grew in all mice (**Figure 1I**). Furthermore, tumor mass, which was measured at endpoint for each individual mouse, was not different between diet groups and suggests that tumor burden was the driver of survival endpoint in both DIO and control groups (**Figure 1I**). These data demonstrate that our organoid orthotopic transplantation model recapitulates key features of obesity-associated colon cancer biology.

**Figure 1.**
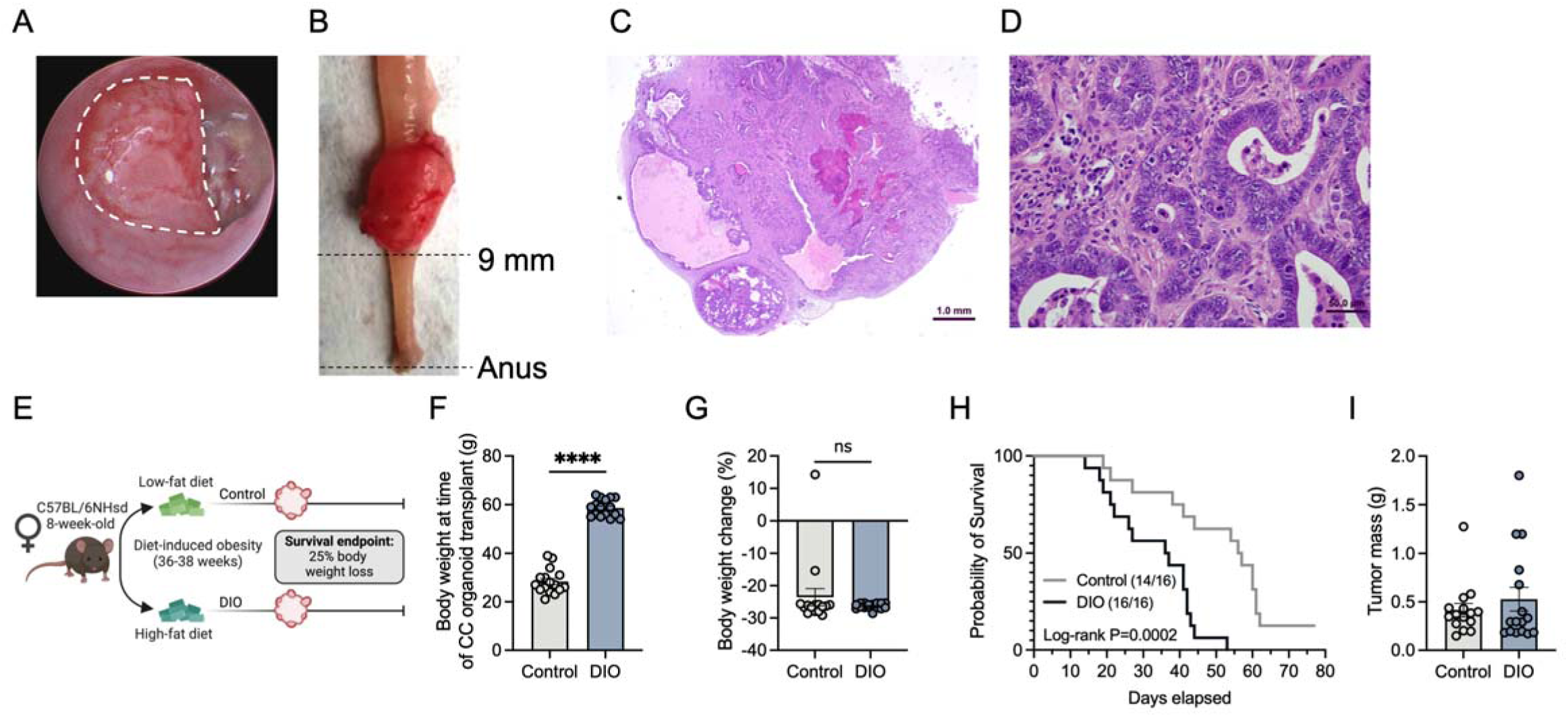
Obesity accelerates morbidity in colon tumor-bearing mice. **(A)** Representative colonoscopy image of tumor growing in the submucosal layer of the colonic wall. Hash marks outline the tumor. **(B)** Representative image of a tumor at time of euthanasia. Representative images of hematoxylin and eosin staining of **(C)** a tumor section and **(D)** glands, diverse immune cell types, and fibrotic material. **(E)** Survival study design. **(F)** Body weight at time of orthotopic injection of AKPST organoids (n=16/16). **(G)** Percent change in body weight from day of organoid injection to euthanasia (n=16/16). **(H)** Kaplan-Meier survival curve. Survival was determined as the number of days after tumor organoid injection that a tumor-bearing mouse remained at or under 25% body weight loss (n=16/16). **(I)** Tumor mass measured at endpoint for each mouse (n=16/16). Data represented as mean ± SEM (F-G, I). Statistical significance determined by unpaired two-sided Student’s t-test (F-G, I) or Mantel-Cox test (H).

### Obesity remodels metabolic and molecular profiles in an orthotopic murine organoid model of colon cancer

To identify metabolic changes potentially driving obesity-related tumor progression, we orthotopically transplanted AKPST organoids into a separate cohort of Control (non-obese) and DIO mice (following >19 weeks of diet treatment) and terminated the study at an early time point (4 weeks) when tumor burden was not significantly different between groups. This approach allowed for the assessment of changes associated primarily with differences in body weight phenotype rather than tumor burden (**Supplemental Figure 1A**). Terminal body weights and mesenteric VAT mass were increased in DIO, relative to Control, mice (**Supplemental Figure 1B-C**). To delineate how obesity shapes tumor cell transcriptomic profiles, we performed bulk RNA-seq analysis on the epithelial (EpCAM+) tumor cell fraction from Control and DIO mice.

Principal component analysis of the top 500 most differentially expressed genes in EpCAM+ cells revealed significant separation of the Control and DIO groups (**Figure 2A**). Gene set enrichment analysis (GSEA) using EpCAM+ RNAseq transcriptomic data and the Molecular Signatures Database (MSigDB) Hallmarks gene sets revealed stark enrichment of proliferation, metabolism, and inflammation-related gene sets in EpCAM+ tumor cells isolated from DIO mice relative to Controls (**Figure 2B**). Within these Hallmark gene sets, multiple genes were redundantly contained within the leading edges of these distinct pathways and processes (**Figure 2C-E**).

**Figure 2.**
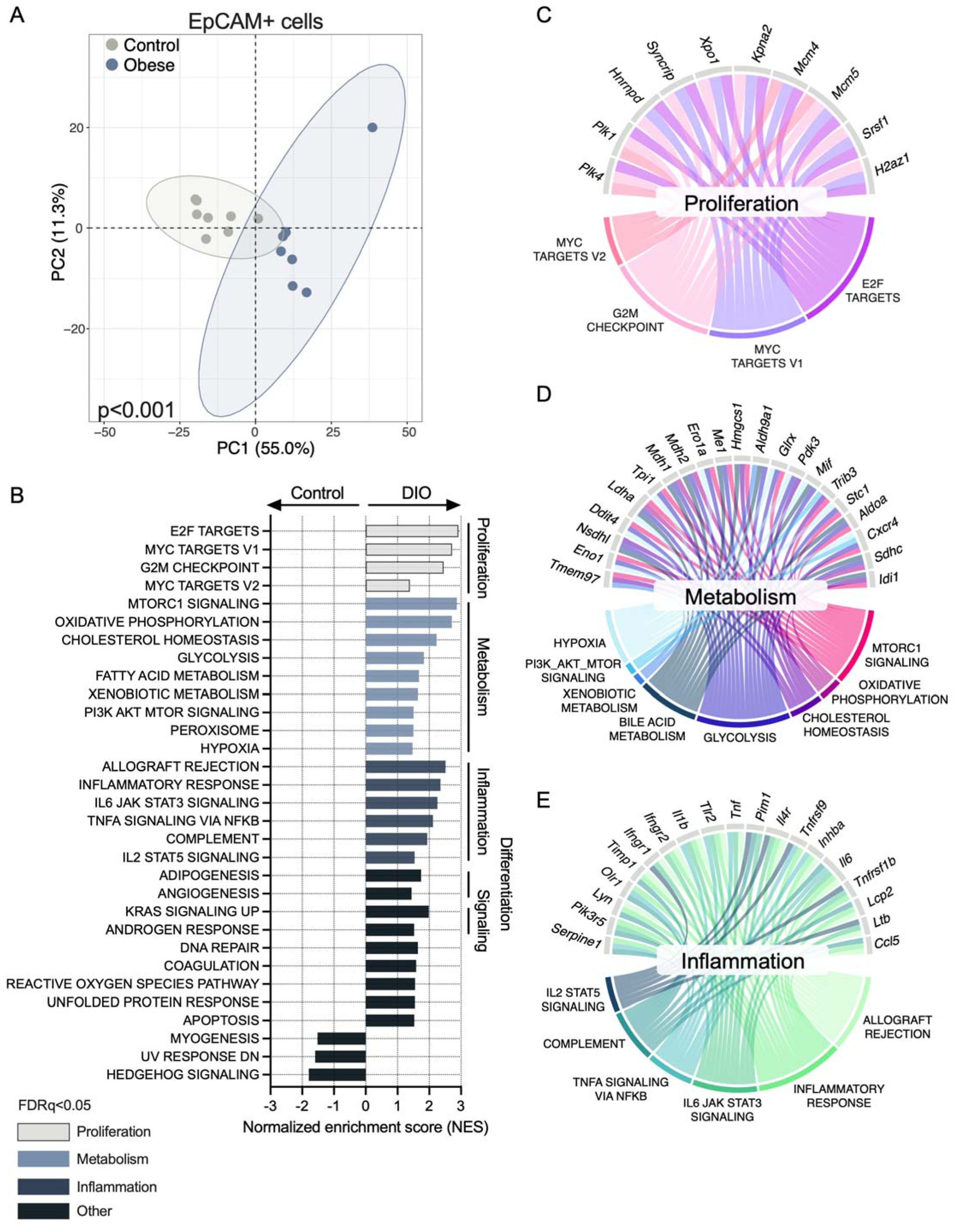
Obesity promotes metabolic and inflammatory processes in EpCAM+ colon tumor cells. **(A)** Principal component analysis of the top 500 most differentially expressed genes. **(B)** Gene set enrichment analysis (GSEA) using EpCAM+ RNAseq transcriptomic data and the MSigDB Hallmarks gene sets. Chord diagrams illustrating which genes contained within the leading edge of the indicated gene sets belong to at least **(C)** 3 proliferation-related, **(D)** 3 metabolism-related, or **(E)** 3 inflammation-related Hallmark gene sets.

### Inflammatory and metabolic pathways are concordantly enriched by obesity in human and mouse colon tumors

We next sought to determine whether the marked transcriptional changes identified in the tumors of obese mice were consistent in human colon tumors. We obtained 193 colon tumors from patients diagnosed with stage I-III colon cancer who are naïve to neoadjuvant treatment from the ColoCare Study, a unique multi-site international prospective survivorship cohort (**Table 1**) (21). Patients were classified as being normoweight (body mass index [BMI] <25 kg/m^2^; 22.7% of study population), overweight (BMI ≥25-30 kg/m^2^; 41.9% of study population) or obese (BMI ≥30 kg/m^2^; 35.5% of study population) (**Table 1**).

GSEA of transcriptomic profiling on patient-derived tumor samples revealed that, similar to our findings in the murine colon cancer model, a set of metabolism and inflammation pathways were uniformly enriched in tumors from patients with obesity vs. normoweight (**Figure 3A**). Moreover, within metabolism- and inflammation-related gene sets, the specific gene sets HYPOXIA, XENOBIOTIC METABOLISM, FATTY ACID METABOLISM, COMPLEMENT, ALLOGRAFT REJECTION, INFLAMMATORY RESPONSE, and IL6 JAK3 STAT3 SIGNALING were all concordantly enriched by obesity in tumors from both mice and humans. While stratifying patient groups as obese (BMI ≥30 kg/m^2^) or normoweight (<25 kg/m^2^) resulted in the greatest overlap between mouse and human gene sets, there remained a consistent enrichment and substantial murine-human overlap of metabolism- and inflammation-related gene sets even when comparing either obese vs. non-obese (BMI <30 kg/m^2^) or overweight+obese (BMI ≥25 kg/m^2^) vs. normoweight (**Supplemental Figure 2A-B**). In contrast to findings in the murine model, proliferation processes (MYC TARGETS V2, G2M CHECKPOINT, MYC TARGETS V1, AND E2F TARGETS) were uniformly suppressed in tumors from patients with obesity vs. normoweight, indicating that obesity exerts differential, species-specific effects on proliferation pathways in the colon tumor (**Figure 3A**).

**Figure 3.**
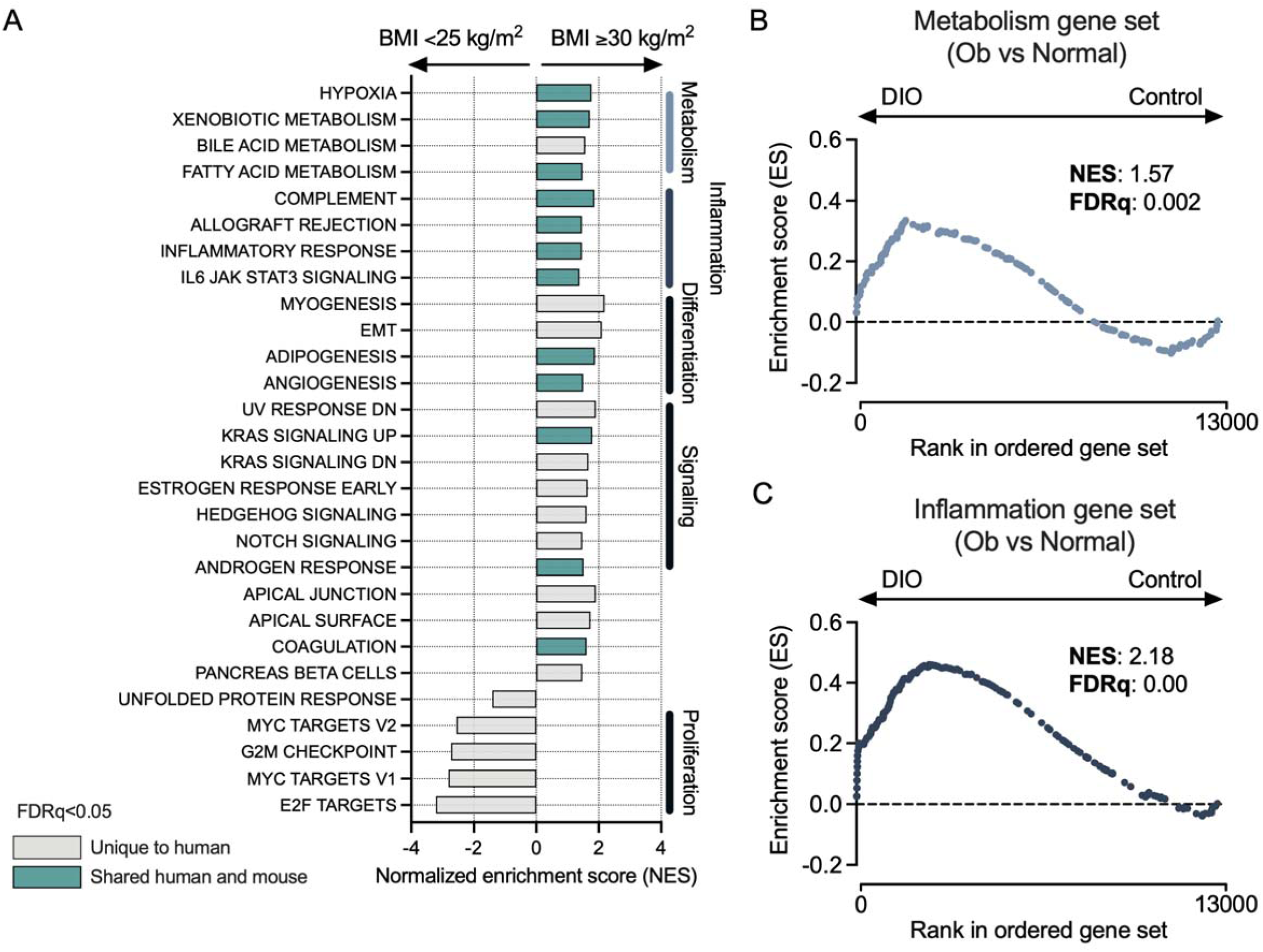
**Obesity-associated enrichment of metabolic- and inflammation-related pathways is conserved between human and mouse colon tumors**. **(A)** Gene set enrichment analysis (GSEA) using global transcriptomic data from human colon tumors. Groups were stratified by patient BMI (<25 kg/m^2^ or ≥30 kg/m^2^). Hallmark gene sets commonly enriched in both human and mouse datasets are color coded in green. Hallmark gene sets unique to human colon tumors are color coded in grey. **(B)** GSEA enrichment plot of mouse EpCAM+ colon tumor RNA-seq data using a custom metabolism gene set. This gene set was constructed from the leading edge genes shared across all significantly enriched metabolism-related pathways identified in the human tumor dataset when stratifying patients by obese vs. normal weight. **(C)** GSEA enrichment plot of mouse EpCAM+ colon tumor RNA-seq data using a custom inflammation gene set. This gene set was constructed from the leading edge genes shared across all significantly enriched inflammation-related pathways identified in the human tumor dataset when stratifying patients by obese vs. normal weight.

Given the robust overlap of specific gene sets across our murine and human analyses, we next examined if strong enrichment scores of enriched gene sets were driven by similar genes in colon tumors from mice and humans. To do so, we created custom gene sets comprised of the leading edge genes from significantly enriched metabolism- and inflammation-related gene sets across the previously defined BMI comparisons of our patient population.

These gene sets reflected the sum of all genes that drove obesity-associated enrichment of metabolism- and inflammation-related gene sets in human colon cancer samples. To determine whether obesity similarly remodeled transcriptomic profiles in mice and human colon tumors, we performed GSEA on our murine EpCAM+ RNAseq transcriptomic data using our custom gene sets derived from patient-derived data. (**Figure 3B-C, Supplemental Figure 2C-F**). For both comparisons of obese vs. normoweight and obese+overweight vs. normoweight, the custom metabolism gene sets were significantly enriched in EpCAM+ tumor cells from obese mice (**Figure 3B, Supplemental Figure 2C-D**). All three custom inflammation gene sets showed significant enrichment in the EpCAM+ population from obese mice, with normalized enrichment scores higher than metabolism gene sets, suggesting a stronger conservation of obesity-driven inflammation-related genes than metabolism-related genes (**Figure 3C, Supplemental Figure 2E-F**). The leading edges from the two custom gene sets presented in Figure 3B-C contained 119 unique genes, representing the overlap between metabolism- and inflammation-related leading edge genes in obesity-associated human and mouse colon cancer (**Supplemental Table 1**). Several of these genes are related to processes involved in lipid metabolism (e.g., *PLIN2*, *ACAT2*, *CD36*, *PPDRD*), innate immune recognition (e.g., *TLR2, MYD88, IRF4)*, and tumor microenvironment modulation (e.g., *MMP9, MMP13, TGFB1, SERPINE1*).

### VAT-derived ligands promote inflammation-related pathways in obesity-driven colon cancer

To ascertain whether dysregulated VAT contributes to the obesity-associated metabolic and/or inflammatory processes observed in colon tumors through tissue-to-tissue signaling, we performed transcriptomic analyses of 188 paired human tumor-adjacent mesenteric VAT and tumor samples. To assess VAT-to-tumor signals, we utilized the FANTOM 5 (30), CellPhoneDB (31), and CellChat (32) databases to identify ligands in the VAT transcriptomic dataset and their intended receptors in the tumor transcriptomics dataset (**Figure 4A**). Using a correlation cutoff >0.3, we identified 88 VAT ligand-tumor receptor pairs (123 nonredundant genes) across the three databases that exhibited robust positive correlation only in VAT-tumor pairs from patients with BMI >30 kg/m^2^ (**Figure 4B; Supplemental Table 2**). Using these 88 ligand-receptor pairs, we performed over-representation analysis with EnrichR (33) to identify significantly enriched MSigDB Hallmark 2020 gene sets (**Figure 4C**). As anticipated when investigating ligand-receptor interactions, many signaling-related gene sets, including UV RESPONSE DOWNREGULATED and WNT BETA CATENIN SIGNALING, were over-represented across the concordantly upregulated 88 VAT ligand-tumor receptor pairs (**Figure 4C**). Moreover, four inflammation-related gene sets, INFLAMMATORY RESPONSE, ALLOGRAFT REJECTION, IL6 JAK STAT3 SIGNALING, and IL2 STAT5 SIGNALING, were consistently enriched in the over- representation analysis (using human VAT-tumor pairs) and in at least one GSEA (using human tumors) pairwise comparison (**Figure 4D**). These results indicate that obesity-associated colon cancer is associated with VAT-to-tumor signaling that enriches for inflammation-related pathways.

**Figure 4.**
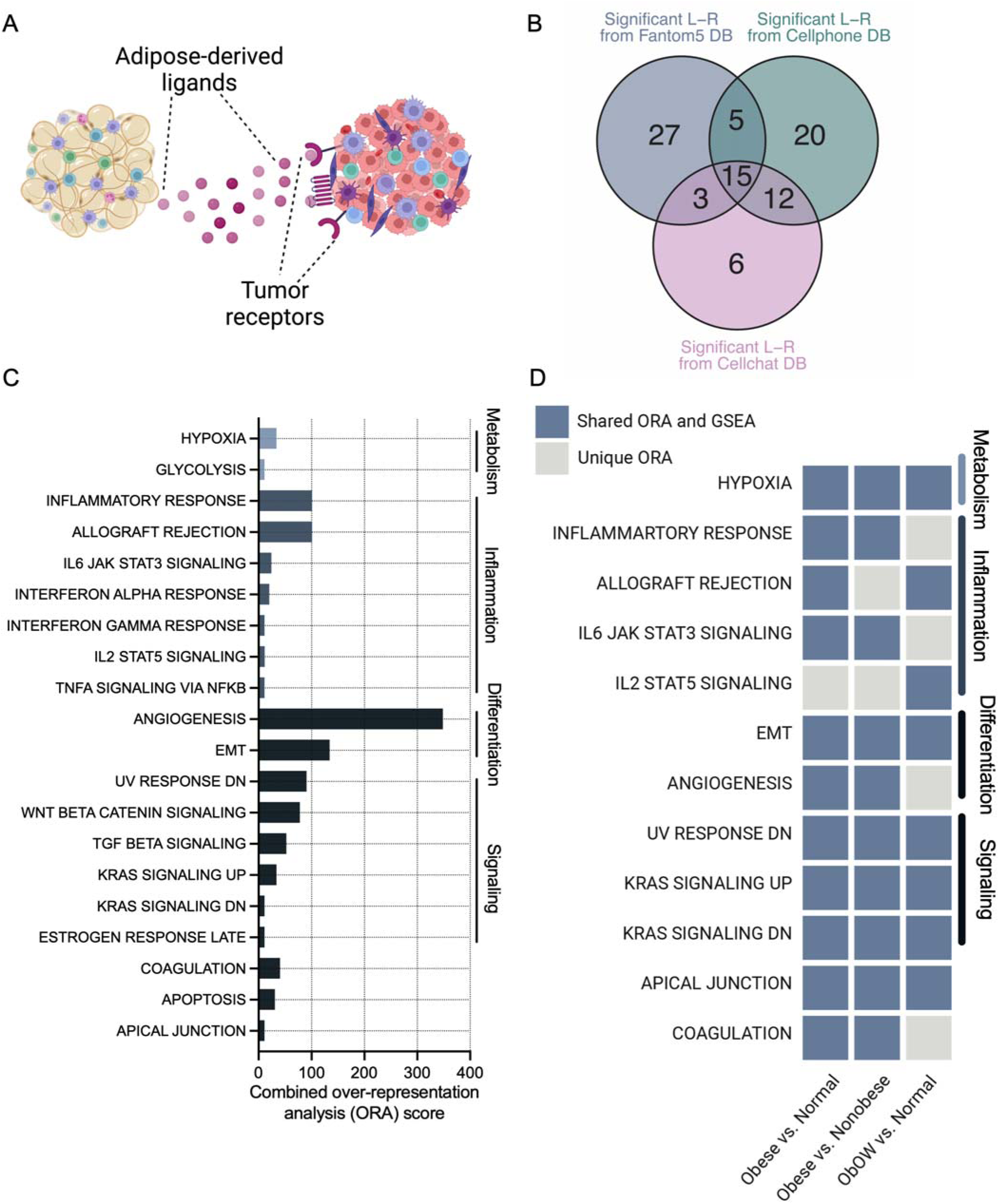
Ligand-receptor analysis reveals VAT-tumor interactions that enrich for inflammation-related pathways in obesity. **(A)** Schematic depicting VAT-derived ligands and tumor-expressed receptors for the ligand-receptor analysis. **(B)** Venn diagram displaying number of ligand-receptor interactions uniquely identified in paired VAT-tumor samples from patients with obesity (BMI ≥30 kg/m^2^). Threshold for significance was defined as Pearson’s correlation >0.3 between ligand and receptor expression using three established ligand-receptor databases: FANTOM 5, CellPhoneDB, and CellChat. **(C)** Over-representation analysis identifying enriched MSigDB Hallmark gene sets using the pre-defined list of significant ligand-receptor interactions uniquely identified in paired VAT-tumor samples from patients with obesity. **(D)** Heatmap indicating which MSigDB Hallmark gene sets were concordantly enriched in both the over-representation analysis and GSEA analysis or were unique to the over-representation analysis. Groups stratified by BMI relate to separate GSEA analyses.

## DISCUSSION

We report inflammation-related and, to a more modest degree, metabolic-related transcriptomic signatures as important, clinically relevant foci of VAT-to-tumor signals that contribute to colon cancer in obesity. Aberrant metabolic and inflammatory signaling are key regulators of colon cancer development and progression (34, 35). Attempts to disentangle inflammatory and metabolic cues originating from different cell types comprising the tumor microenvironment, including infiltrating immune cells and tumor cells, are challenging (36). Therefore, to identify changes in gene expression specific to tumor cells, we aligned the transcriptomic profiles of bulk-sequenced isolated EpCAM+ murine tumor cells and human colon cancer samples. We found that obesity promoted consistent enrichment of inflammation- and metabolic-related signatures across species.

We also established key signatures of obesity-driven transcriptomic remodeling of the colon tumor environment, which are highly consistent at even modest elevations of BMI above 25 kg/m^2^. Considerable work has attempted to understand if and how BMI corresponds to metabolic impairment, and whether superior measures of metabolic health could be used in place of, or in addition to, BMI to appropriately stratify risk for chronic diseases like colon cancer (37–39). Our key findings were robust across three distinct BMI thresholds (obese vs. normoweight, obese+overweight vs. normoweight, and obese vs. non-obese), each of which were associated with the inflammatory and metabolic tumoral signaling networks that we identified. However, we are unable to determine whether alternative measures of metabolic disease or increased adiposity similarly remodel colon cancer transcriptomic programs. Indeed, colon tumors from patients with type 1 diabetes versus type 2 diabetes demonstrate distinct gene expression profiles (40). Finally, we have previously shown that the amount of VAT is correlated with distinct transcriptomic patterns in VAT (9). Thus, our results indicate that while stratification by BMI is sufficient to identify colon tumor remodeling, contributions from other factors, such as the amount of visceral fat and specific metabolic dysfunction are possible.

Using transcriptomic profiling, we identified that the inflammation-related leading edge genes overlapping between murine and human transcriptomic datasets were related to innate immune sensing (*TLR2*, *MYD88*, and *IRF4*) and tumor microenvironment remodeling (*MMP9*, *TGFB1*, *SERPINE1*). Production of immunomodulatory proteins such as TGFβ and SERPINE1, which we found to be associated with obesity across mouse and human colon cancer, engenders an immunosuppressive tumor microenvironment and thereby promotes colon cancer progression (41–44). TGFβ production in colon tumors strongly predicts disease progression, and inhibition of TGFβ signaling not only restores tumor immunosurveillance but also limits tumor progression (41, 42). Importantly, such restoration of immunosurveillance is potently synergistic with immune checkpoint inhibition(42). SERPINE1, like TGFβ, is abundantly expressed by colon tumors (43), and enhances colon tumor growth through blunting of antitumor immunity, specifically by promoting the exclusion of cytotoxic CD8+ T cells from the tumor microenvironment (44). Obesity promotes TGFβ and SERPINE1 expression in contexts outside of colon cancer (45, 46); thus, this signaling axis may be translationally actionable in limiting obesity-driven colon cancer.

In regard to the obesity-related changes in immune-sensing genes, toll-like receptor (TLR) proteins sense pathogen- or damage-associated molecular patterns to induce innate immune signaling responses and play important roles in the development of various inflammatory and metabolic diseases (47). In humans and murine models, these pathways are critical for the maintenance of effective intestinal barrier function and physical separation from the intestinal microbiota (48). MYD88, an adaptor protein critical for a range of TLR signaling, is required for intestinal barrier function (48). Epithelial cell intrinsic TLR2-MYD88 signaling supports the proliferation of tumor cells and enables carcinogenesis (49). Similarly, MYD88 dependent TLR4 signaling supports the acceleration of colon cancer progression by high-fat diet feeding (50). High-fat diets themselves remodel the intestinal microbiome and promote the presence of microbes that produce metabolites that can impair intestinal barrier function and thereby promote colon cancer carcinogenesis (51). By identifying, for example, TLR2-MYD88 signaling as a conserved feature of obesity-induced inflammation in colon cancer, this work bridges critical mechanistic findings on the role of inflammation in cancer progression with the modifiable risk factor of obesity.

In addition to synthesizing preclinical and clinical findings by integrating transcriptomic signatures across two species, another strength of our study was the assessment of putative interactions between VAT-derived factors and tumor signaling in clinical samples. It is well-established that VAT expands and becomes dysregulated in obesity and is a poor prognostic factor for colon cancer, highlighting the potential role of adipose-derived cytokines, adipokines, lipid-mediators and other factors to promote the incidence and progression of colon cancer (8, 9, 11). Specifically in colon cancer, mesenteric VAT envelopes the colonic serosa and our clinical team collected VAT samples within 1-3 cm of the tumor to ensure anatomical proximity to the cancer. Using three established databases to build an extensive list of ligand-receptor pairs, we found that correlated expression (Pearson’s coefficient >0.3) of cognate pairs of ligands from VAT and receptors in tumors from patients with obesity were overrepresented across 7 inflammation-related gene sets (INFLAMMATORY RESPONSE, ALLOGRAFT REJECTION, IL6 JAK STAT3 SIGNALING, INTERFERON ALPHA RESPONSE, INTERFERON GAMMA RESPONSE, IL2 STAT5 SIGNALING, TNFA SIGNALING VIA NFKB). Thus, specifically in patients with obesity, signals from mesenteric VAT may be one source driving inflammatory responses in colon tumors and further work elucidating these signaling networks may be tractable in orthotopic murine models.

Obesity remodels metabolic programs of various tissues at both cellular and systemic levels, and our aligned preclinical and clinical datasets identify metabolic alterations that are conserved between obesity-driven colon cancer in mice and humans. Specifically, we found numerous lipid metabolism genes to be induced by obesity in colon cancer (e.g., *PLIN2*, *ACAT2*, *CD36*, and *PPDRD*). Colon tumors enhance the expression of lipogenic enzymes to sustain considerable enrichment of TAG relative to normal tissue (52). Yet the roles of dietary lipid supply and obesity in supporting such metabolic profiles are poorly understood. Our finding that enrichment of lipid metabolism-related genes is a feature of obesity-driven transcriptomic remodeling in colon cancer cells demonstrates that while lipid metabolism is reprogrammed by tumor cell-intrinsic processes, colon cancer cells remain sensitive to the systemic metabolic state.

In summary, integrating complementary data from well-controlled mouse studies and clinical samples from a representative sample of patients with newly diagnosed colon cancer, we identified obesity-responsive, cancer-promoting pathways and genes common to mice and humans. These findings while informative, cannot replace the need for definitive mechanistic studies to define the extent to which these pathways direct colon cancer progression. In our murine organoid model of colon cancer, obesity reduced survival and enriched several MSigDB Hallmark gene sets involving inflammation, metabolism, and proliferation. In patients with colon cancer, obesity enriched inflammation and metabolic pathways in concordance with our mouse model findings. Moreover, human transcriptomic analyses also uncovered procancer interactions between VAT-derived ligands and tumor-intrinsic receptors. Synthesis of our murine and human findings enables prioritization of new molecular targets underlying the obesity-colon cancer association and provides a strong foundation for future translational studies to develop and test mechanism-based lifestyle or pharmacologic interventions.

## ACKNOWLEDGEMENTS

The authors thank all participating patients and acknowledge assistance from the Biorepository and Molecular Pathology and the High-Throughput Genomics and Cancer Bioinformatics Shared Resources at the HCI at the University of Utah (supported in part by the National Cancer Institute of the National Institutes of Health under award number P30CA042014). The research was facilitated by the Moffitt Cancer Center’s Total Cancer Care Protocol and the Tissue Core Shared Resource at the H. Lee Moffitt Cancer Center & Research Institute, an NCI-designated Comprehensive Cancer Center (P30CA076292). We also thank the NCT Biobank at the Heidelberg University Hospital. We are grateful to Dr. Jeffrey Everitt for histopathology review of the AKPST tumors, and Dr. Chris Stubben for his useful input regarding the bioinformatic analyses. Illustrations were created in BioRender.com.

## AUTHOR CONTRIBUTIONS

*Designed research*: E.M.G., T.L., V.M.B., A.C.T., J.R., C.M.U., S.D.H.; *Conducted research*: E.M.G., T.L., B.M., S.S.K., B.G., C.A.W., O.A., M.F.C., A.C., C.B.; P.S., I.S., S.H., J.N.C., J.J., A.B.; *Administered project*: V.M.B., C.A.W., C.H., J.O., C.M.U.; *Provided essential materials*: B.G., M.A.S., C.K., E.M.S., D.A.B., A.T.T., D.S., C.I.L., J.C.F., A.C.T., J.R., C.M.U., S.D.H.; *Analyzed data*: E.M.G., T.L., M.F.C. D.A.N., K.B.; *Wrote paper*: E.M.G., S.D.H. All authors have read and approved the final manuscript.

## DATA AVAILABILITY

Murine transcriptomic data have been deposited to GEO with accession number GSE284066 and will be publicly available as of the date of publication. ColoCare Study data are available from on reasonable request and as described on the ColoCare website (https://uofuhealth.utah.edu/huntsman/labs/colocare-consortium/). Our data sharing procedures are available online

(https://uofuhealth.utah.edu/huntsman/labs/colocare-consortium/data-sharing/new-projects.php). For any additional questions please contact the ColoCare Study Administrator Team. Further information and requests for resources and reagents should be directed to and will be fulfilled by the lead contact Stephen D. Hursting.

---

**Supplemental Figure 1.** O**b**esity **drives metabolic dysfunction in tumor-bearing mice. (A)** AKPST tumor weights (n=13/8), **(B)** body weight (n=13/8), and **(C)** mesenteric fat mass (n=12/8) in C57BL/6Hsd mice after 4 weeks of orthotopic tumor growth. Data represented as mean ± SEM. Statistical significance determined by unpaired two-sided Student’s t-test (A-C).

**Supplemental Figure 2.** E**l**evated **BMI is associated with metabolic- and inflammation-related pathways that are conserved between human and mouse colon tumors. (A-B)** Gene set enrichment analysis (GSEA) using global transcriptomic data from human colon tumors. Hallmark gene sets commonly enriched in both human and mouse datasets are color coded in green. Hallmark gene sets unique to human colon tumors are color coded in grey. Groups were stratified by **(A)** patient BMI <30 kg/m^2^ vs. ≥30 kg/m^2^ and **(B)** patient BMI <25 kg/m^2^ vs. ≥25 kg/m^2^. **(C-F)** GSEA of EpCAM+ colon tumor cell RNA-seq data from DIO and Control mice using custom sets constructed from the leading edge genes shared across all significantly-enriched metabolism- or inflammation-related pathways identified in the human tumor datasets. GSEA enrichment plot of the metabolism gene set created when stratifying patients by **(C)** obese (Ob) vs. non-obese (NoOb) or **(D)** obese+overweight (ObOW) vs. normal weight. GSEA enrichment plot of mouse EpCAM+ colon tumor RNA-seq data using custom inflammation gene sets. GSEA enrichment plot of the inflammation gene set created when stratifying patients by **(E)** obese (Ob) vs. non-obese (NoOb) or **(F)** obese+overweight (ObOW) vs. normal weight.

