## Supplementary figures and images for "Obesity promotes conserved inflammatory and metabolic transcriptional programs in mouse and human colon tumors"

### Supplemental figure 1

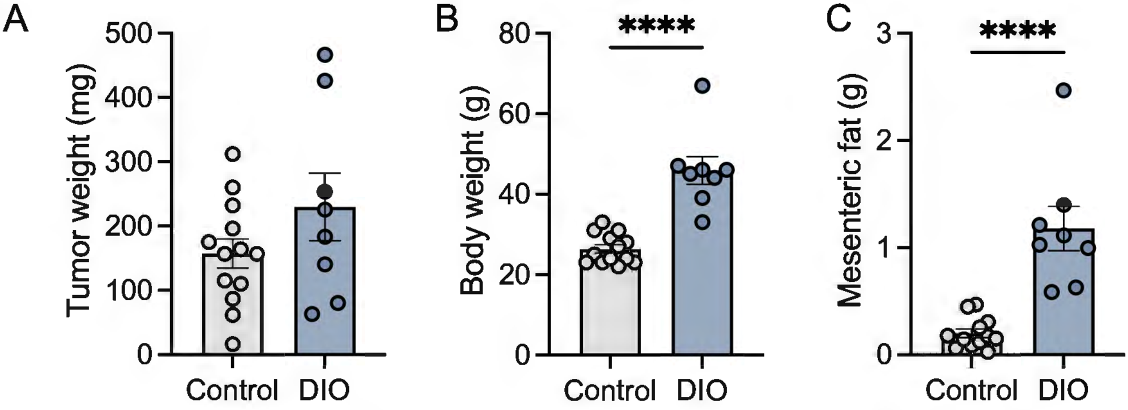

### Supplemental figure 2

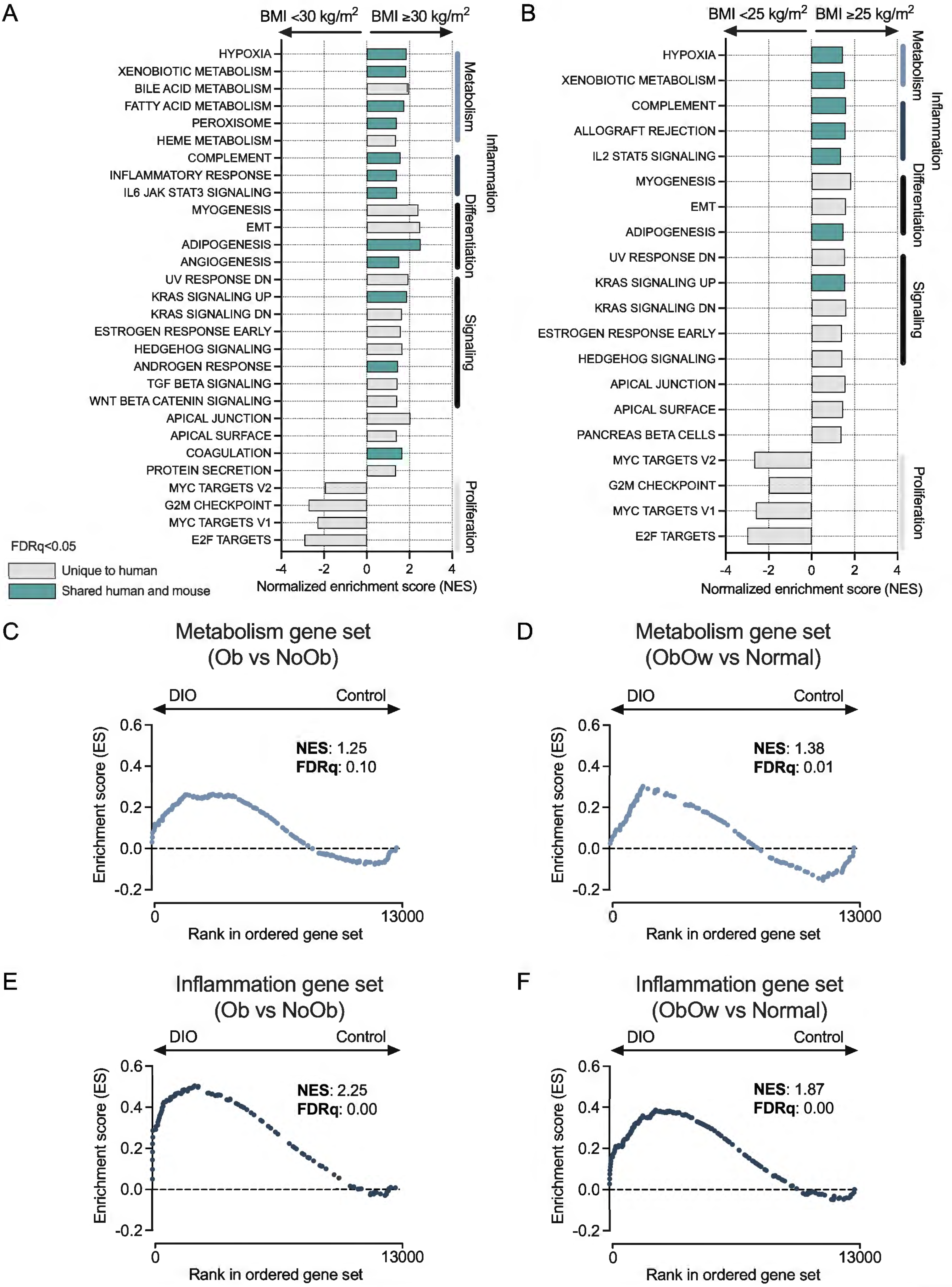
