## Supplemental table 1 for "Obesity promotes conserved inflammatory and metabolic transcriptional programs in mouse and human colon tumors"

**Supplemental Table 1.** Metabolism- and inflammation-leading edge genes shared amongst DIO mice and patients with obesity.

| METABOLISM | | INFLAMMATION | |
|---|---|---|---|
| Gene symbol | Rank in EpCAM+ gene list | Gene symbol | Rank in EpCAM+ gene list |
| *G0S2* | 20 | *Sell* | 14 |
| *Lgals1* | 39 | *Gpr132* | 16 |
| *Pdk4* | 48 | *Mmp9* | 19 |
| *Plin2* | 130 | *Il10Ra* | 24 |
| *Sult2B1* | 165 | *Fgr* | 29 |
| *Rragd* | 170 | *Fcer1G* | 35 |
| *Ets1* | 268 | *Ccr7* | 40 |
| *Ddit4* | 310 | *Spi1* | 51 |
| *Pnrc1* | 363 | *Hla-Dob* | 84 |
| *Serpine1* | 364 | *Inhbb* | 86 |
| *Pdk3* | 417 | *Ets1* | 268 |
| *Igfbp3* | 553 | *Tgfb1* | 287 |
| *Hmox1* | 564 | *Msrb1* | 335 |
| *Acads* | 582 | *Serpine1* | 364 |
| *Cda* | 708 | *Ptafr* | 436 |
| *Pdgfb* | 759 | *Ctsd* | 476 |
| *Ninj1* | 842 | *Cntfr* | 509 |
| *Acox2* | 882 | *Tlr2* | 516 |
| *Bhlhe40* | 886 | *Itgb3* | 551 |
| *Entpd5* | 900 | *Hmox1* | 564 |
| *Cxcr4* | 945 | *Ltb* | 671 |
| *Slc25A1* | 1034 | *Cda* | 708 |
| *Cyfip2* | 1046 | *Pfn1* | 713 |
| *Glrx* | 1077 | *Pdgfb* | 759 |
| *Stbd1* | 1092 | *Cfp* | 785 |
| *F10* | 1127 | *Ptprc* | 808 |
| *Btg1* | 1150 | *Itga5* | 888 |
| *Cat* | 1210 | *Thy1* | 939 |
| *Chst2* | 1266 | *Emp3* | 956 |
| *Man1A1* | 1275 | *Mmp13* | 959 |
| *Ndrg1* | 1360 | *Ccr2* | 972 |
| Angptl4 | 1454 | *St8Sia4* | 1013 |
| *Cavin3* | 1455 | *Hcls1* | 1052 |
| *Pkp1* | 1460 | *Prkcb* | 1089 |
| *Slc46A3* | 1495 | *Cd86* | 1122 |
| *P4Ha2* | 1578 | *F10* | 1127 |
| *Cd36* | 1602 | *Itga4* | 1165 |
| *Tat* | 1610 | *Cd82* | 1171 |
| *Ppard* | 1630 | *Gng2* | 1192 |
| *Pgf* | 1651 | *Fyn* | 1208 |
| *Gstt2* | 1695 | *Il7R* | 1217 |
| *Ccng2* | 1735 | *Irf4* | 1219 |
| *Cited2* | 1769 | *Chst2* | 1266 |

| METABOLISM | | INFLAMMATION | |
|---|---|---|---|
| Gene symbol | Rank in EpCAM+ gene list | Gene symbol | Rank in EpCAM+ gene list |
| | | *Rhog* | 1285 |
| | | *Ctsb* | 1309 |
| | | *Map4K1* | 1315 |
| | | *Cr2* | 1370 |
| | | *Bak1* | 1408 |
| | | *Gata3* | 1410 |
| | | *Was* | 1512 |
| | | *Dock10* | 1517 |
| | | *Hdac9* | 1543 |
| | | *C3Ar1* | 1552 |
| | | *Itgam* | 1599 |
| | | *Cd36* | 1602 |
| | | *Rasgrp1* | 1622 |
| | | *Ccnd3* | 1644 |
| | | *Fpr1* | 1674 |
| | | *Fcgr2B* | 1684 |
| | | *Itgal* | 1737 |
| | | *Gpr183* | 1857 |
| | | *Csf2Ra* | 1861 |
| | | *Kcnj2* | 1900 |
| | | *Cybb* | 1925 |
| | | *Gnb2* | 1944 |
| | | *Ctss* | 1949 |
| | | *Ccl20* | 1993 |
| | | *Ly86* | 2016 |
| | | *Il16* | 2044 |
| | | *Il18R1* | 2146 |
| | | *Cd79A* | 2169 |
| | | *Gnai2* | 2185 |
| | | *Cdk5R1* | 2209 |
| | | *Ifngr2* | 2225 |
| | | *Fyb1* | 2277 |
| | | *Ccl22* | 2288 |
| | | *Cmklr1* | 2320 |
| | | *Slc7A2* | 2343 |
| | | *Pik3R5* | 2404 |
| | | *Notch4* | 2416 |
| | | *Pdpn* | 2537 |
| | | *C1Qc* | 2577 |
| | | *Myd88* | 2581 |
| | | *Il10Rb* | 2587 |
