## Supplemental table 2 for "Obesity promotes conserved inflammatory and metabolic transcriptional programs in mouse and human colon tumors"

**Supplemental Table 2.** Significant (Pearson's correlation >0.3) ligand-receptor pairs unique to VAT-tumor pairs from patients with BMI >30 kg/$m^2$. VAT ligand-tumor receptor pairs were identified using 3 databases: FANTOM 5, CellPhoneDB, and CellChat.

| Ligand-Receptor pair | Ligand gene symbol | Receptor gene symbol | Database(s) |
|---|---|---|---|
| *ANGPTL1_ITGB1* | *ANGPTL1* | *ITGB1* | CellChat |
| *APOA1_ABCA1* | *APOA1* | *ABCA1* | CellPhoneDB |
| *APOA1_CUBN* | *APOA1* | *CUBN* | CellPhoneDB |
| *B2M_KIR2DL1* | *B2M* | *KIR2DL1* | FANTOM 5 |
| *CCL17_CCR4* | *CCL17* | *CCR4* | FANTOM 5 / CellPhoneDB / CellChat |
| *CCL17_CCR8* | *CCL17* | *CCR8* | FANTOM 5 |
| *CD5L_CD5* | *CD5L* | *CD5* | FANTOM 5 |
| *CDH2_FCER2* | *CDH2* | *FCER2* | CellPhoneDB |
| *CORT_SSTR5* | *CORT* | *SSTR5* | FANTOM 5 / CellPhoneDB / CellChat |
| *CXCL12_ITGB1* | *CXCL12* | *ITGB1* | FANTOM 5 |
| *CXCL13_CXCR3* | *CXCL13* | *CXCR3* | FANTOM 5 / CellChat |
| *DCN_MET* | *DCN* | *MET* | FANTOM 5 |
| *DDC_DRD4* | *DDC* | *DRD4* | CellPhoneDB |
| *DDC_HTR1B* | *DDC* | *HTR1B* | CellPhoneDB |
| *DMP1_ITGAV* | *DMP1* | *ITGAV* | CellChat |
| *DPEP2_CYSLTR1* | *DPEP2* | *CYSLTR1* | CellPhoneDB |
| *F2_F2R* | *F2* | *F2R* | FANTOM 5 / CellPhoneDB / CellChat |
| *F2_F2RL3* | *F2* | *F2RL3* | FANTOM 5 / CellPhoneDB / CellChat |
| *F2_ITGAV* | *F2* | *ITGAV* | CellPhoneDB |
| *F2_THBD* | *F2* | *THBD* | FANTOM 5 / CellPhoneDB |
| *FGF2_SDC4* | *FGF2* | *SDC4* | FANTOM 5 |
| *FGF5_FGFR1* | *FGF5* | *FGFR1* | FANTOM 5 / CellPhoneDB / CellChat |
| *FLT3LG_FLT3* | *FLT3LG* | *FLT3* | FANTOM 5 / CellPhoneDB |
| *GAL_GALR3* | *GAL* | *GALR3* | FANTOM 5 / CellPhoneDB / CellChat |
| *GDF11_TGFBR1* | *GDF11* | *TGFBR1* | CellPhoneDB / CellChat |
| *GGT1_CYSLTR2* | *GGT1* | *CYSLTR2* | CellPhoneDB |
| *GLS2_GRIN2A* | *GLS2* | *GRIN2A* | CellPhoneDB |
| *GLS2_GRM5* | *GLS2* | *GRM5* | CellPhoneDB |
| *GNAI2_CAV1* | *GNAI2* | *CAV1* | FANTOM 5 |
| *GNAI2_EDNRA* | *GNAI2* | *EDNRA* | FANTOM 5 |
| *GNAI2_F2R* | *GNAI2* | *F2R* | FANTOM 5 |
| *GNAI2_OPRM1* | *GNAI2* | *OPRM1* | FANTOM 5 |
| *GNAS_GLP1R* | *GNAS* | *GLP1R* | FANTOM 5 |
| *HLA-B_KIR2DL3* | *HLA-B* | *KIR2DL3* | FANTOM 5 |
| *HLA-C_KIR2DL1* | *HLA-C* | *KIR2DL1* | FANTOM 5 / CellPhoneDB / CellChat |
| *HLA-C_KIR2DL3* | *HLA-C* | *KIR2DL3* | FANTOM 5 / CellPhoneDB / CellChat |
| *HLA-C_KIR2DS4* | *HLA-C* | *KIR2DS4* | FANTOM 5 / CellChat |
| *IL12B_IL12RB2* | *IL12B* | *IL12RB2* | FANTOM 5 / CellPhoneDB / CellChat |
| *IL19_IL20RB* | *IL19* | *IL20RB* | FANTOM 5 / CellPhoneDB / CellChat |
| *INHBC_ACVR2A* | *INHBC* | *ACVR2A* | FANTOM 5 / CellPhoneDB / CellChat |
| *JAM2_F11R* | *JAM2* | *F11R* | CellChat |
| *L1CAM_ITGA5* | *L1CAM* | *ITGA5* | FANTOM 5 / CellPhoneDB |
| *LTA_CACHD1* | *LTA* | *CACHD1* | FANTOM 5 |
| *MICB_HCST* | *MICB* | *HCST* | CellPhoneDB |
| *MICB_KLRK1* | *MICB* | *KLRK1* | CellPhoneDB |
| *MMP7_ERBB4* | *MMP7* | *ERBB4* | FANTOM 5 |
| *NGF_MAGED1* | *NGF* | *MAGED1* | FANTOM 5 |

| Ligand-Receptor pair | Ligand gene symbol | Receptor gene symbol | Database(s) |
|---|---|---|---|
| *NRG4_ERBB2* | *NRG4* | *ERBB2* | FANTOM 5 / CellChat |
| *NTNG2_LRRC4C* | *NTNG2* | *LRRC4C* | FANTOM 5 |
| *PENK_OGFR* | *PENK* | *OGFR* | FANTOM 5 |
| *PF4_CXCR3* | *PF4* | *CXCR3* | FANTOM 5 / CellPhoneDB / CellChat |
| *PF4_LDLR* | *PF4* | *LDLR* | FANTOM 5 |
| *PF4_SDC2* | *PF4* | *SDC2* | FANTOM 5 |
| *PF4_THBD* | *PF4* | *THBD* | FANTOM 5 |
| *PF4V1_CXCR3* | *PF4V1* | *CXCR3* | CellChat |
| *PNOC_OPRL1* | *PNOC* | *OPRL1* | FANTOM 5 / CellPhoneDB |
| *RIMS2_ABCA1* | *RIMS2* | *ABCA1* | FANTOM 5 |
| *RSPO1_LRP6* | *RSPO1* | *LRP6* | FANTOM 5 / CellPhoneDB |
| *RSPO3_SDC4* | *RSPO3* | *SDC4* | FANTOM 5 |
| *SLC10A4_CHRM2* | *SLC10A4* | *CHRM2* | CellPhoneDB |
| *SLC18A1_DRD4* | *SLC18A1* | *DRD4* | CellPhoneDB |
| *SLC1A2_GRM1* | *SLC1A2* | *GRM1* | CellPhoneDB |
| *SLC1A7_GRM5* | *SLC1A7* | *GRM5* | CellPhoneDB |
| *SLC5A7_CHRM1* | *SLC5A7* | *CHRM1* | CellPhoneDB |
| *SLC6A12_GABRA3* | *SLC6A12* | *GABRA3* | CellPhoneDB |
| *SLC6A2_ADRA1D* | *SLC6A2* | *ADRA1D* | CellPhoneDB |
| *TGFB1_CAV1* | *TGFB1* | *CAV1* | FANTOM 5 |
| *TGFB1_CD109* | *TGFB1* | *CD109* | FANTOM 5 |
| *TGFB1_CXCR4* | *TGFB1* | *CXCR4* | FANTOM 5 |
| *TGFB1_ENG* | *TGFB1* | *ENG* | FANTOM 5 |
| *TGFB1_TGFBR1* | *TGFB1* | *TGFBR1* | FANTOM 5 / CellPhoneDB / CellChat |
| *TGFB3_TGFBR1* | *TGFB3* | *TGFBR1* | FANTOM 5 / CellPhoneDB / CellChat |
| *TNN_SDC4* | *TNN* | *SDC4* | CellChat |
| *TPH1_HTR1B* | *TPH1* | *HTR1B* | CellPhoneDB |
| *VTN_ITGA3* | *VTN* | *ITGA3* | FANTOM 5 |
| *WNT1_FZD1* | *WNT1* | *FZD1* | FANTOM 5 / CellPhoneDB / CellChat |
| *WNT1_FZD2* | *WNT1* | *FZD2* | CellPhoneDB / CellChat |
| *WNT10A_FZD1* | *WNT10A* | *FZD1* | CellPhoneDB / CellChat |
| *WNT10A_FZD4* | *WNT10A* | *FZD4* | CellChat |
| *WNT16_FZD1* | *WNT16* | *FZD1* | CellPhoneDB / CellChat |
| *WNT16_FZD10* | *WNT16* | *FZD10* | CellPhoneDB / CellChat |
| *WNT16_FZD2* | *WNT16* | *FZD2* | CellPhoneDB / CellChat |
| *WNT16_FZD4* | *WNT16* | *FZD4* | CellPhoneDB / CellChat |
| *WNT7A_FZD1* | *WNT7A* | *FZD1* | CellPhoneDB / CellChat |
| *WNT7A_FZD2* | *WNT7A* | *FZD2* | CellPhoneDB / CellChat |
| *WNT7A_FZD4* | *WNT7A* | *FZD4* | CellPhoneDB / CellChat |
| *WNT7A_FZD8* | *WNT7A* | *FZD8* | CellPhoneDB / CellChat |
| *WNT7B_FZD2* | *WNT7B* | *FZD2* | CellPhoneDB / CellChat |
